# Spatially resolved single-cell atlas reveals cortical glial reactivity associated with regions of neuronal growth programs in both hemispheres after stroke

**DOI:** 10.64898/2026.10.26.761917

**Authors:** Matthew Kenwood, Dene Betz, Eduardo Guttierez-Kuri, Bryan Derebery, Priscilla Rhoton, Juan Pablo Palavicini, Denise M. O. Ramirez, Mark P. Goldberg

**Affiliations:** Department of Neurology, University of Texas at San Antonio; Graduate School of Biomedical Science, University of Texas at San Antonio; Sam and Ann Barshop Institute for Longevity and Aging Studies, University of Texas Health Science Center San Antonio (University of Texas at San Antonio); Department of Cellular and Integrative Physiology, University of Texas at San Antonio; Department of Medicine, Diabetes Division, University of Texas at San Antonio; Department of Neurology, University of Texas Southwestern Medical Center; Peter O’Donnell, Jr. Brain Institute, University of Texas Southwestern Medical Center

## Abstract

Recovery after focal motor cortex stroke relies on structural remodeling of neuronal circuits in the perilesional cortex (PLC) and contralesional cortex (CLC). Although transcriptomic studies have identified distinct reactive glial states after stroke, how their spatial organization relates to processes of tissue degeneration and neuronal circuit remodeling remains poorly understood. We used a custom 347-gene Xenium spatial transcriptomics panel to profile approximately 650,000 cells across both hemispheres of male and female mice 7 days after photothrombotic stroke. We identified injury-associated myeloid populations concentrated within the infarct and reactive astrocytes enriched in the glial scar. Distinct reactive populations also extended into spared cortex, including an interferon-associated myeloid population and a reactive astrocyte population that was present in both PLC and CLC. Spatial mapping further revealed elevated neuronal growth-associated gene expression in both regions, particularly among layer 5 extratelencephalic projection neurons, which include corticospinal neurons. Intriguingly, reactive glial populations were enriched in regions exhibiting elevated neuronal growth-associated gene expression, including areas of the CLC containing axons and presynaptic terminals originating from the PLC. Together, these findings reveal a striking spatial association between reactive glial populations and neurons exhibiting elevated growth-associated gene expression and support glial-neuronal interactions as potential targets for promoting circuit remodeling and functional recovery after stroke.

## Introduction

Structural remodeling after motor cortical stroke occurs in spared neuronal circuits in both the perilesional (PLC)^1–4^ and contralesional cortex (CLC)^5–9^ where it drives functional recovery. ^10–12^ Neurons in both regions undergo axonal sprouting and synaptic reorganization, which occurs along with the activation of growth-associated transcriptional programs.^7,13,14^ Non-neuronal cells, including astrocytes and microglia, can influence these processes through inflammatory signaling, extracellular matrix remodeling, and trophic support. Although distinct reactive glial states have been identified after stroke, their functional roles remain incompletely understood.^15^ Defining how these states are distributed throughout the post-stroke brain, including relative to the infarct, damaged white matter tracts, and spared cortical regions may provide insight into their biological roles and potential contributions to post-stroke circuit remodeling.

Transcriptomic studies have identified diverse reactive glial states associated with injury and repair after stroke.^16–19^ However, single-cell RNA sequencing lacks spatial information^20–22^, while previous spatial transcriptomic studies have been limited by multicellular resolution or a focus on the injured hemisphere.^21,23^ Our recent identification of shared inflammatory transcriptional signatures in the PLC and CLC further raises the question of whether specific reactive glial states extend into spared regions and how their distribution relates to neurons actively undergoing remodeling.^19^ To address this, we used a custom 347-gene Xenium spatial transcriptomics panel to profile approximately 650,000 cells across both hemispheres of male and female mice 7 days after cortical stroke. We mapped reactive glial populations in relation to lesion anatomy, neuronal growth-associated gene expression, and PLC cortical circuitry.

## Results

### Single-Cell Spatial Transcriptomics Reveals Regional Heterogeneity After Stroke

We performed unilateral photothrombotic stroke or sham surgery targeting the sensorimotor cortex in male and female adult C57BL6/J mice (n=4 stroke, n=2 sham). T2-weighted MRI at 24 hours confirmed cortical infarcts averaging 22.6 ± 0.74 mm³ (Figure 1A, , Sup 1A). Seven days after stroke, Xenium spatial transcriptomics yielded approximately 650,000 cells across the PLC, CLC, and subcortical structures, with approximately 94% of detected transcripts assigned to cells after Proseg segmentation (Figure 1A-C). ^22^

**Figure 1:**
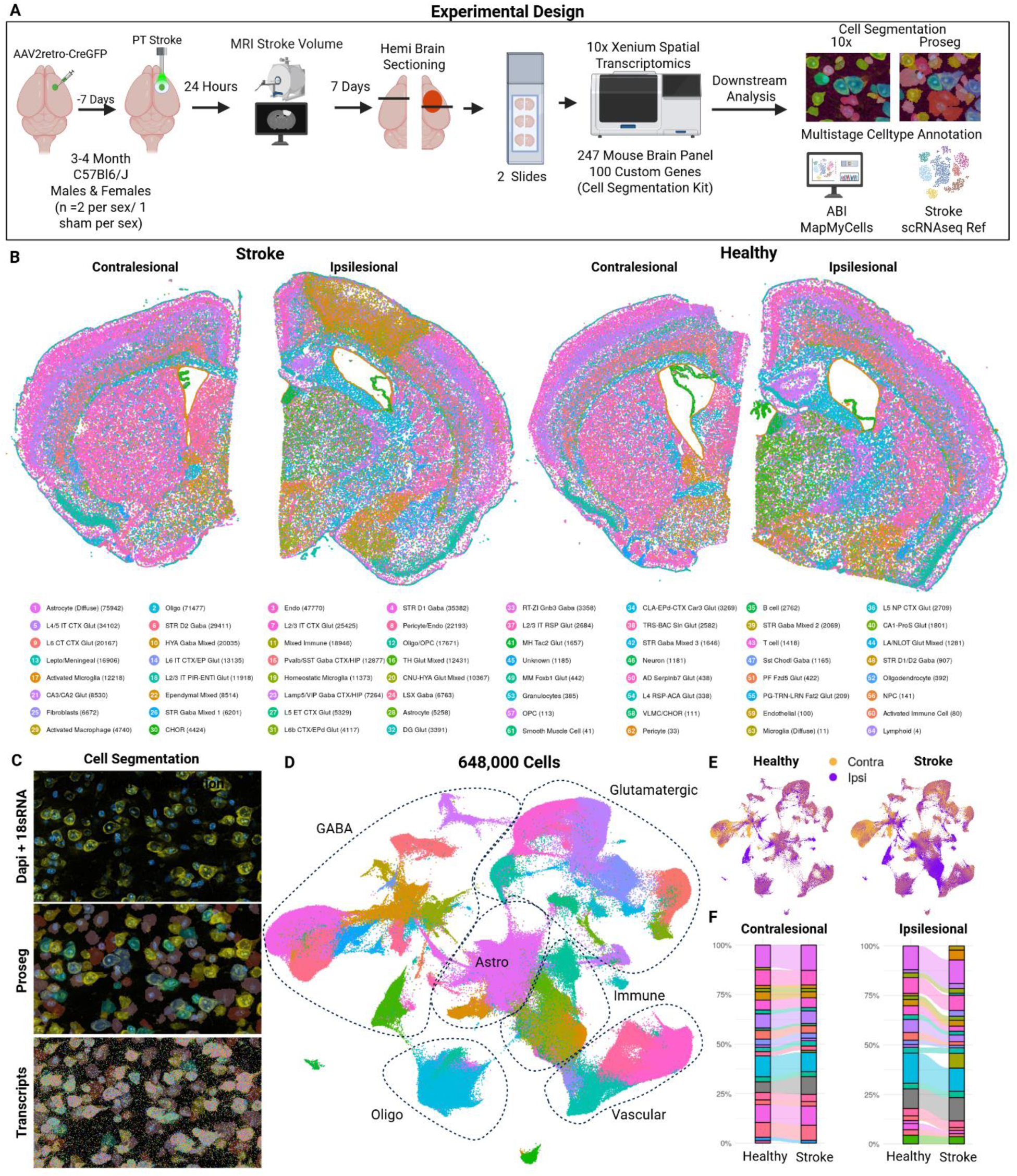
Spatial Transcriptomics of Cortical Model of Stroke during the Subacute Phase. (A) Experimental timeline of 10x Xenium experiment (B). Contralesional and Ipsilesional hemi-sections of segmented cells in coordinate space of female stroke and healthy controls colored by cell type manually curated subclass annotation (C) Example cell segmentation outline of 10x cell segmentation kit staining (top), proseg voxel segmentation (middle), and segmentation and all transcripts (bottom) from xenium explorer (D) UMAP of 625,888 cells across all conditions colored by subclass and outlined by higher order cell types.(E) UMAPs split by stroke condition and colored by contralesional and ipsilesional hemisphere (F) Alluvial proportion plots of stroke conditions and hemisphere colored by cell type annotations of cells with > 1% of total proportions. Cell colors are uniform from panels B & C.

Stroke produced pronounced changes in the cellular composition of the injured hemisphere (Figure 1D–F). Neuronal loss was confined to the infarct and spared neurons were detected immediately beyond the glial scar. The infarct contained diverse glial, immune, and stromal populations, including fibroblasts and infiltrating immune cells concentrated in the lesion core (Sup 2). Activated and infiltrating myeloid cells exhibited the greatest increased proportions, comprising approximately 10% of all cells profiled in PLC sections. In contrast, the CLC showed no substantial changes in overall cell-type proportions (Figure 1F). Together, these findings highlight differences in cellular responses to stroke across cortical regions associated with post-stroke remodeling.

### Spatial Organization of Reactive Myeloid Populations After Stroke

Given the substantial changes in myeloid cell proportions following stroke, we next characterized their transcriptional profiles and spatial distribution. We reclustered myeloid cells using BANKSY^23^, a spatial clustering method that incorporates each cell’s transcriptional profile and gene expression from its surrounding cellular neighborhood to identify populations associated with distinct tissue environments (Figure 2A). Myeloid cells segregated into three broad groups: *Siglech*⁺/*Spp1*⁻ cells characteristic of resident microglia, *Siglech*⁻/*Spp1*^low^ cells, and *Siglech*⁻/*Spp1*^high^ cells associated with injury-induced myeloid activation (Figure 2A; Sup 3A–C).

**Figure 2:**
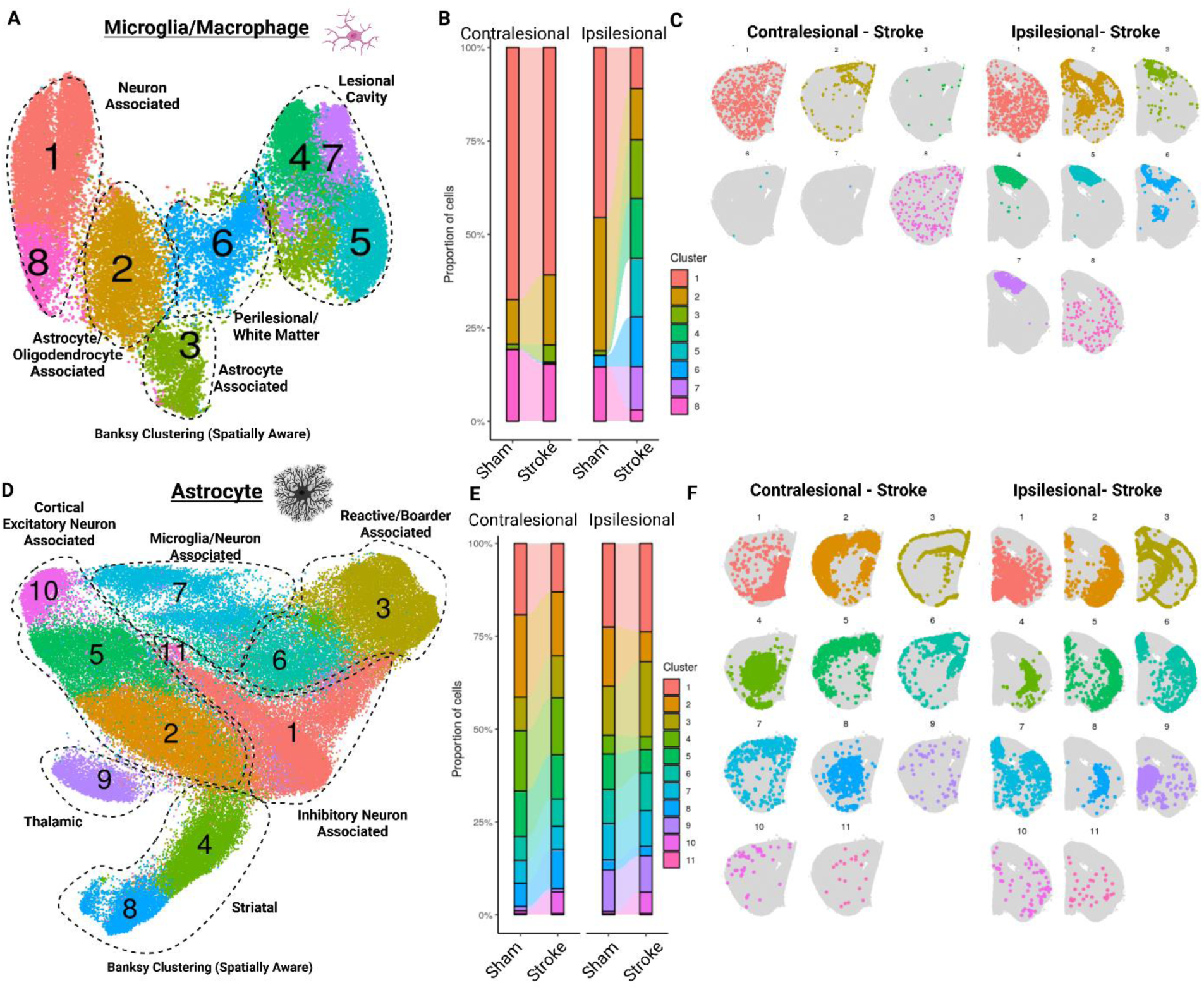
Myeloid and Astrocyte Responses to Localized Cortical Stroke are Not Confined to Ipsilesional Hemisphere: (A) UMAP of microglia and macrophages (Myeloid cells) clustered by BANKSY neighborhood clustering (B) Proportion alluvial plots of myeloid BANKSY clusters between sham and stroke samples split by PLC and CLC (C) Spatial Coordinate plots of myeloid BANKSY clusters split by PLC and CLC (D) UMAP of astrocytes clustered by BANKSY neighborhood clustering (E) Proportion alluvial plots of astrocyte BANKSY clusters between sham and stroke samples split by PLC and CLC (F) Spatial Coordinate plots of astrocyte BANKSY clusters split by PLC and CLC

*Siglech*⁺/*Spp1*⁻ myeloid cells (cluster 1) were diffusely distributed throughout the brain and associated with neuronal markers, including *Caln1*, *Necab2*, and *Calb1*. Other populations showed greater regional specificity. *Siglech*⁻/*Spp1*^low^ populations (clusters 4, 5, and 7) were concentrated within fibroblast-rich regions of the infarct. In contrast, *Siglech*⁻/*Spp1*^high^ reactive populations (clusters 2 and 6) occupied cellular environments enriched for astrocyte (*Gfap*), oligodendrocyte (*Opalin*), oligodendrocyte precursor cell (*Gpr17*), and T-cell (*Trbc2*) markers (Sup 3C).

Cluster 6 exhibited a pronounced injury-associated signature, with increased *Spp1*, *Igf1*, and *Cd44*, resembling previously described reactive microglial populations^24^ (Supplemental Figure 3F). This population was localized to ipsilesional hemisphere, including PLC, striatum, corpus callosum, and internal capsule. Cluster 2 occupied similar regions in the ipsilesional hemisphere but was also present in the contralesional corpus callosum and cortex (Figure 2B,C). Compared with homeostatic populations (clusters 1 and 8), cluster 2 showed increased *Igfbp5*, *Ly6a*, and *Slfn5*, consistent with interferon-associated signaling response^24^.Together, these findings distinguish injury-associated myeloid populations and populations that localize to spared regions in both ipsilesional and remote contralesional hemispheres.

### Spatial Organization of Reactive Astrocyte Populations After Stroke

Given the interactions between astrocytes and myeloid cells after stroke, we next used BANKSY to characterize astrocyte populations and their cellular neighborhoods. BANKSY identified 11 astrocyte clusters, all of which were detected in both the PLC and CLC, although their proportions differed by region and stroke condition (Figure 2D–F; Sup 4C,D).

Astrocyte clusters were more broadly distributed across both hemispheres compared to myeloid populations. Three clusters localized to anatomical regions including striatum (4, 8) and thalamus (9). The remaining occupied cell-type specific neighborhoods including cortical excitatory neurons (2, 5, and 10), inhibitory neurons (1 and 11), microglia (7), and border-associated glia (3,6) (Sup 4A,B).

Clusters 3 and 6 differed in the magnitude and anatomical distribution of their responses to stroke. Cluster 3 was enriched in the ipsilesional glial scar and exhibited a strong signature consistent with previously reported reactive astrocyte phenotypes^25,26^, with increased *Gfap*, *Stat3*, *Cd44*, and *Laptm5*, and *Cd68* expression, which likely reflected neighboring myeloid cells (Sup 4E). In contrast, cluster 6 showed a more limited transcriptional response, including increased *Laptm5* and *Col1a1*, and was present in both the PLC and CLC. Cluster 10 also increased in both PLC and CLC and localized to cortical neighborhoods enriched for the neuronal growth-associated gene *Sprr1a*, which was significantly upregulated relative to sham (log₂FC > 3; Sup 4E). These findings suggest that reactive astrocyte populations exhibit spatial organization at both regional and cell-type-specific levels.

### Spatial Association of Neuronal Growth-Associated Programs and Reactive Glial Populations

The significant increase in neuronal *Sprr1a* expression raised the question of whether neuronal growth-associated transcriptional programs were localized to specific regions within PLC and CLC. We therefore examined the spatial distribution of growth-associated gene expression across cortical projection neurons using a literature-curated gene set associated with axonal growth and plasticity, including *Gap43*^27^, *Jun*^28^, *Stat3*^29^, *Sprr1a*^30^, *Gdf10*^2^, *Crebbp*^31^, *Bmp4*^32^, *Adcyap1*^33^, *Fos*^34^, *Klf7*^35^, *Klf6*^29^, *Meis2*^36^, *Pbx1*^37^, *Myt1l*^38^, *Smad3*^39^) (Figure 3A). We divided cortical projection neurons into 250-µm spatial bins and mapped the difference in mean growth-associated module scores between stroke and sham mice for three projection neuronal subclasses (L2-6IT, L5 ET, and L6 CT) (Figure 3A).

**Figure 3:**
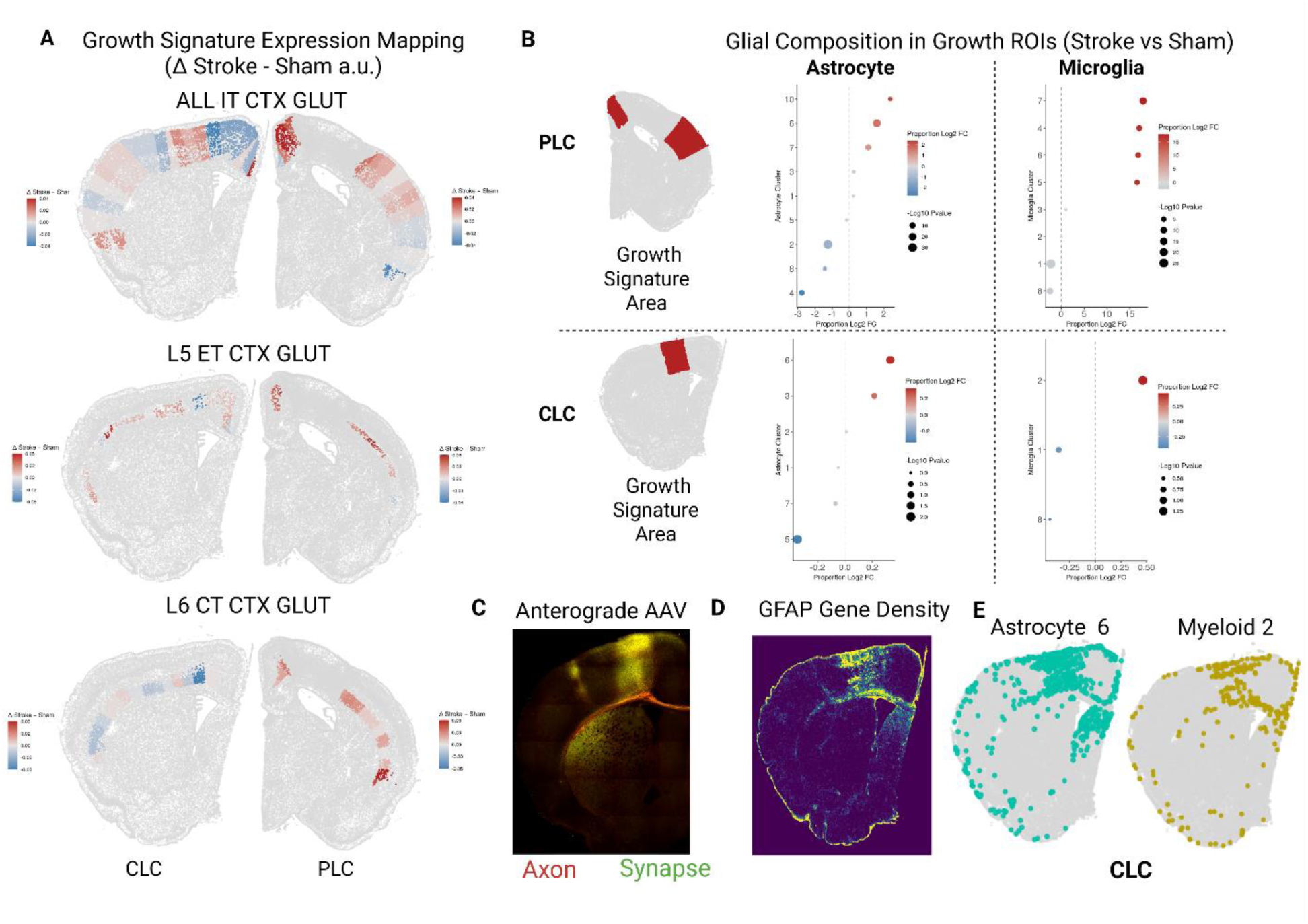
Growth-Associated Gene Expression Mapping of PLC and CLC. (A) Coordinate plots of growth-associated module activation (delta of Stroke – Sham) across 4 neuronal classes: L2/3 IT CTX GLUT (top left), L4/5 IT CTX GLUT (top right), L5 ET CTX GLUT (lower left), and L6 CT CTX GLUT (lower right). Each section is individually scaled; delta mean across all animals is plotted on representative section coordinates from female stroke in CLC and PLC. (B) Cluster proportion log2 fold-change plots of astrocytes and microglia (myeloid) cells inside growth-associated areas detailed in red-shaded regions in the PLC and CLC. (C) STPT images of representative anterograde Synaptotag4 which labels axons (tdTomato), synapses (eGFP) injected into the somatomotor cortex (D) Xenium explorer density plots of a contralesional and ipsilesional representative female section of Gfap expression density (E) Astrocyte cluster 6 and myeloid cluster 2 cell density in the CLC 7d after stroke

Growth-associated module scores were elevated in both the PLC and CLC, although their spatial distribution differed across the projection neuron subclasses. L5 ET neurons, which include corticospinal neurons, showed the most broadly elevated growth-associated scores across both regions, whereas L6 CT neurons exhibited elevated scores only in the PLC (Figure 3A). We next examined whether regions with elevated neuronal growth-associated scores were associated with the glial populations identified in our preceding analyses. Cortical bins exhibiting elevated growth-associated scores were significantly enriched for reactive astrocyte cluster 6 (*Stat3*, *Cd44*, *Laptm5*) in both the PLC and CLC (Figure 3B). Several myeloid populations (clusters 4, 5, 6, and 7) were enriched in growth-associated regions of the PLC, although interferon-associated cluster 2 was the only myeloid population significantly enriched in the growth-associated regions in CLC. Astrocytes in PLC growth-associated regions showed increased *Gfap* and *Col1a1* expression, while myeloid cells in these regions exhibited increased *Spp1*, *Igf1*, *Fn1*, *Col1a1*, *Cd44*, and *Cd68* expression. These changes were not observed in the corresponding CLC regions (Sup 5).

To examine whether these spatial patterns corresponded to cortical connectivity, we performed anterograde tracing using SynaptoTAG4^9^ and serial two-photon tomography to visualize axons and presynaptic terminals originating from the PLC. In the CLC, regions containing labeled projections were enriched for reactive astrocyte cluster 6 and myeloid cluster 2 (Figure 3C–E). These findings support distinct spatial patterns of neuronal growth-associated transcription in the PLC and CLC and suggest that specific reactive glial populations are enriched in regions exhibiting these neuronal programs.

## Discussion

We applied subcellular spatial transcriptomics to resolve the glial and neuronal landscape across PLC and CLC one week after photothrombotic stroke. Two major conclusions arose. First, reactive glial states partition into lesion-restricted and bilateral populations. Scar-associated astrocytes and injury-associated myeloid populations were concentrated within and around the infarct, whereas a reactive myeloid state (cluster 2) and a reactive astrocyte state (cluster 6) extended into the CLC. Although bilateral glial reactivity has been reported by histology and bulk sequencing ^19,40^, our findings distinguish reactive populations extending into spared cortex from those concentrated near the lesion and suggest that the uninjured hemisphere mounts a defined reactive program.

Second, growth-associated transcriptional programs are activated in both the PLC and CLC. Subcerebral projection neurons, positioned to have the greatest effect on motor circuit repair^41^showed the broadest activation of these programs. Structural plasticity of L5 ET neurons after stroke is well documented^6,7,9,42^. Our data show that the transcriptional programs underlying this growth are engaged bilaterally and most broadly in this population versus other cortical projection neurons.

Notably, these neuronal growth programs were concentrated in the same territories that harbored the bilateral reactive glial populations. Gliosis has long been linked to neuronal growth and repair after stroke ^2,43^. Here, we suggest specific glial states contribute to this process in both the PLC and CLC, despite the difference in proximity to the inflammatory environment of the lesion. Because these associations are spatially correlative, we cannot resolve whether reactive glia instruct, permit, or accompany neuronal growth states without targeted perturbation.

Together, these findings reveal distinct spatial patterns of glial reactivity across both hemispheres after stroke, including cortical regions remote from the lesion. The correspondence between reactive glial states and neuronal growth-associated gene expression provides a new framework for investigating how structural plasticity is initiated in spared cortical circuits. Future studies integrating spatial transcriptomics with functional approaches may identify cell-cell signaling pathways that regulate neuronal growth programs and contribute to post-stroke motor recovery.

## Methods

### Animals

C57BL/6J mice (WT) of both sexes were used (12-16 weeks old, 000664 Jackson Laboratory Bar Harbor ME) and maintained on a 12 hour light/dark cycle. All procedures were performed between 9-12am. The Institutional Animal Care and Use Committee of the University of Texas at San Antonio approved all animal procedures.

### Photothrombotic Stroke

Mice were deeply anesthetized with isoflurane and a 45mW 561nm laser is targeted 1.9mm laterally and 0.25mm anterior to bregma to target the right somatomotor cortex. Rose Bengal (40 mg/kg) is injected intraperitoneally. Following 1 minute of dwell, the laser is illuminated for 15 minutes. The scalp is then sutured, disinfected, and extended-release buprenorphine is administered subcutaneously.

### MRI Assessment of Infarct Volumes

MRIs were acquired on a Bruker 7T system with custom coils at 24 hours post-stroke. 128x128 px 1mm coronal slices spanning the whole brain AP. Lesions were quantified by manually tracing hyperintense lesions on T2-weighted images in Fiji-ImageJ.

### Intracranial Injections

One week prior to stroke, mice received a unilateral AAV2r-SYN-creEGFP into the right motor cortex (-1.8 ML/ 0.0 AP) to label neurons that project to the eventual ischemic lesion. For anterograde tracing of the motor cortex, animals underwent identical preparation procedures and were injected with SynaptoTAG4 (AAVDJ-Syn-mTdTomato-Syb2eGFP-WPRE) to label axons with Tdtomato and presynaptic terminals with EGFP.

### Tissue Collection for 10x Xenium

Mice were transcardially perfused with ice-cold RNase-free PBS followed by ice-cold 4% Paraformaldehyde for 5 minutes. Brains were rapidly dissected and placed in 4% PFA for 24-hours followed by 24 hours in 30% sucrose. Brains were then hemisected, embedded separately in OCT blocks, and stored at -80 °C for long-term storage.

### 10x Xenium

A 100 gene panel was built on top of the 10x Mouse Brain Panel version 1.1 using the Xenium panel designer (v4.1.0), totaling 347 genes. We processed Xenium slides using the Fixed Frozen protocol with cell segmentation kit staining. Instrument (v4.0.1.4) and analysis (v4.0.1.0). Xenium Ranger outputs of nuclei segmentation using Dapi were utilized to resegment using Proseg ^24^ (v3.1.1) including reissuing Xenium Ranger (v4.0.1.1).

## Declarations

The authors declare that they have no competing interests.

## Acknowledgements

This work was supported by National Institutes of Health (NIH) awards T32AG082661 (DB), T32GM -113896 and -145432 (DB), T32NS082145-08 (DB), AHA Fellowship 23PRE1018993 (DB), T32NS082145-14 (BD), GSBS Neuroscience Training Fellowship (MK), OT2OD032581 (MPG), R21NS146921-01A1 (MPG) and by gifts from the Haggerty Foundation (MPG) and Meier Family Foundation (MPG). This work was supported by the Alzheimer’s Association AARG-21-846012 (JPP) and the National Institutes of Health (NIH) RL5 scholar P30AG044271 (JPP). Data was generated in the Genome Sequencing Facility, which is supported by UTHSCSA, NIH-NCI P30 CA054174 (Cancer Center at UTHSCSA), NIH Shared Instrument grant 1S10OD030311-01 (S10 grant towards NovaSeq 6000 sequencing system), S10OD038331 (S10 grant to Xenium Analyzer), CPRIT Core Facility Support Award (RP 220662), and NCI R50CA265339 to ZL. Serial two-photon tomography was conducted by the Whole Brain Microscopy Facility at UTSW (RRID: SCR_017949).

---

**Supplemental Figure 1:**
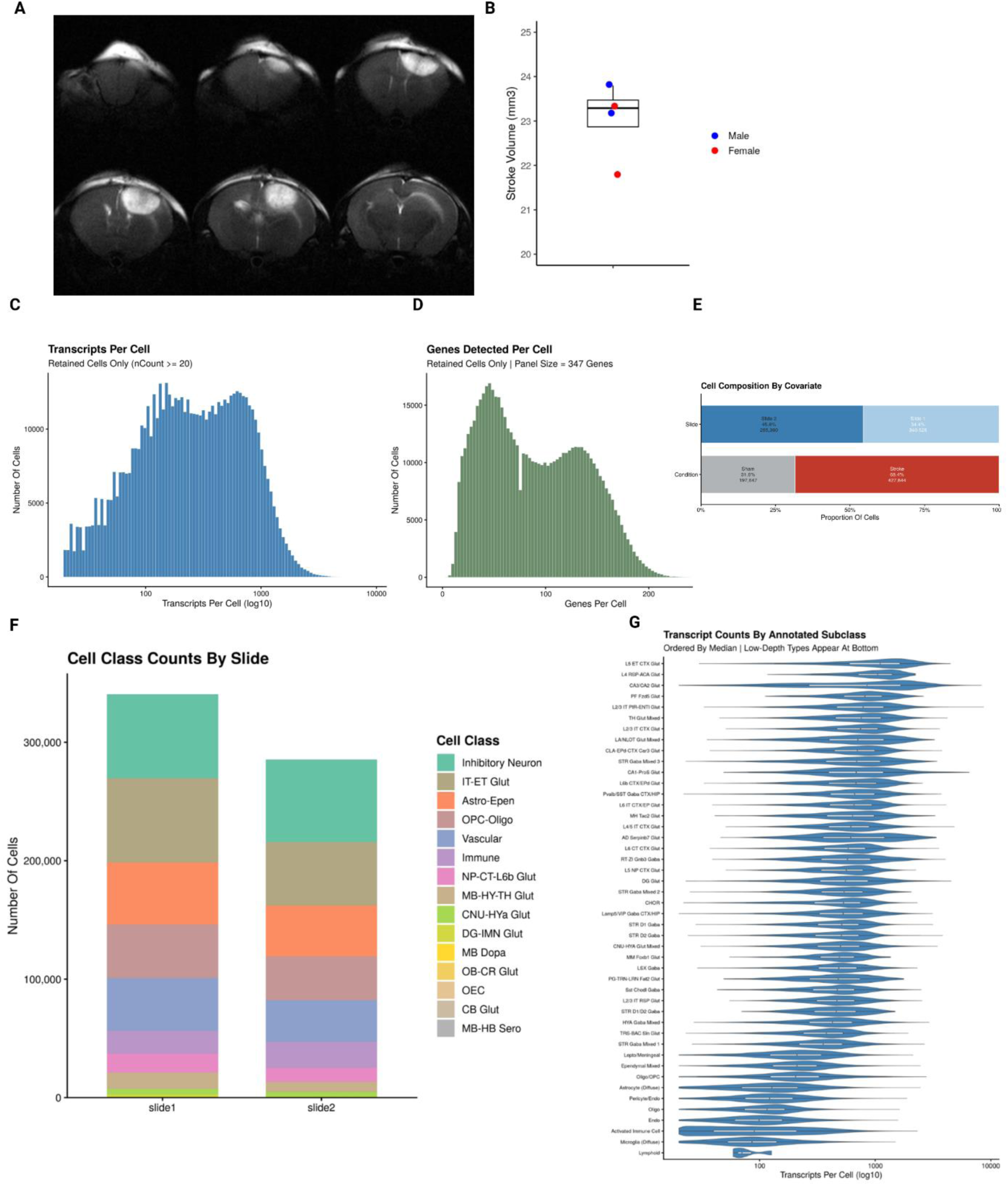
(A) Representative 1 mm T2 MRI images at 24 hours post-stroke (B) Stroke volumes in male and female mice used for Xenium (C) Histogram of transcripts per cell after filtering (D) Histogram of genes detected per cell after filtering (E) Number of cells per slide and by condition (F) Per Xenium slide cell type composition (G) Violin plots of transcript counts per annotated cell types

**Supplemental Figure 2:**
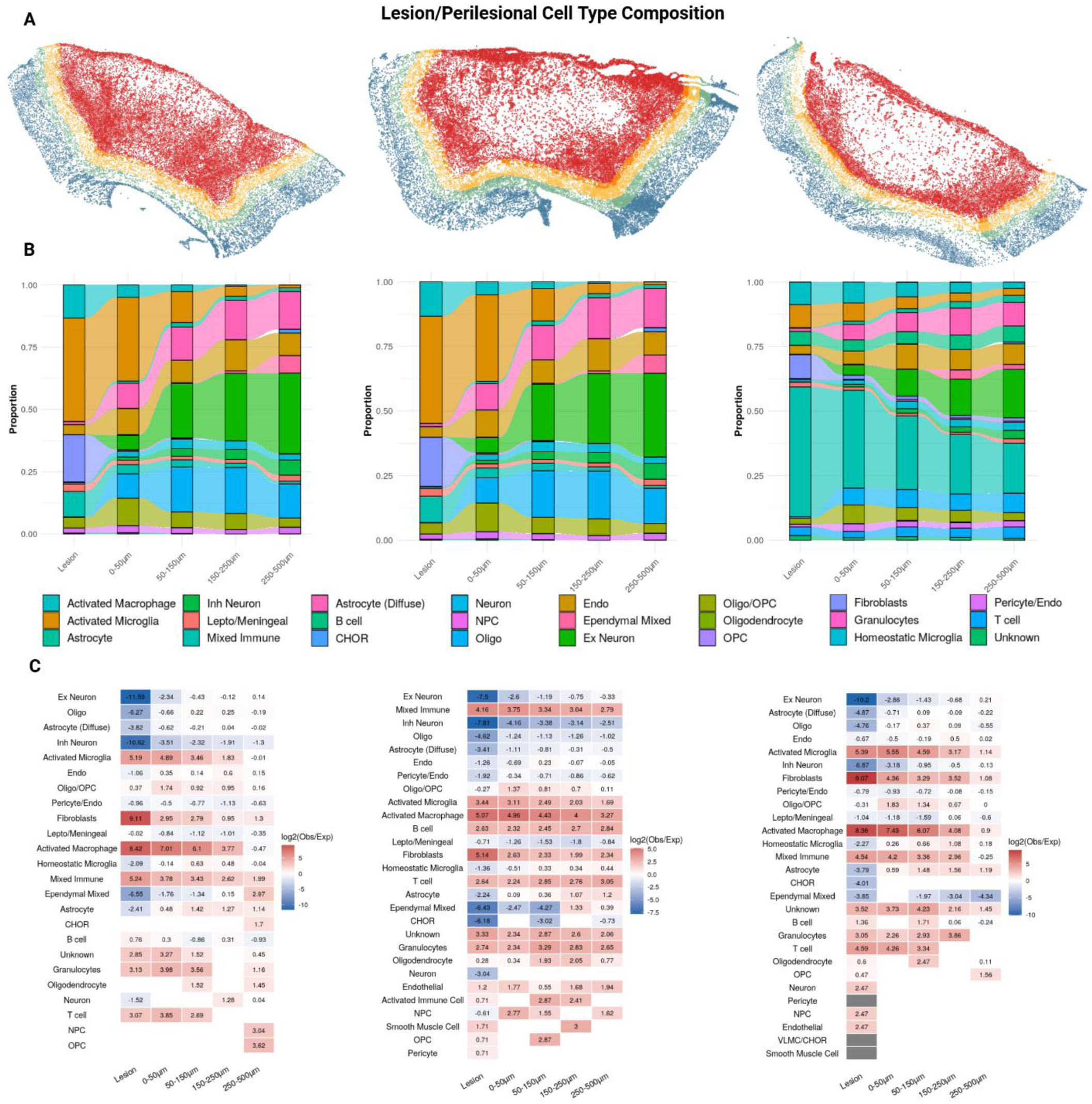
(A) Outlined stroke lesions and cells colored by binned perilesional zones: lesion (red), 0-50um (orange), 50-150um (yellow), 150-250 (teal), 250-500um (blue) (B) Alluvial proportion plots of cell type compositions within the bins of the three lesions (C) Fold change of observed over expected (>500um from lesion) cell type density of each lesion and perilesional bin.

**Supplemental Figure 3:**
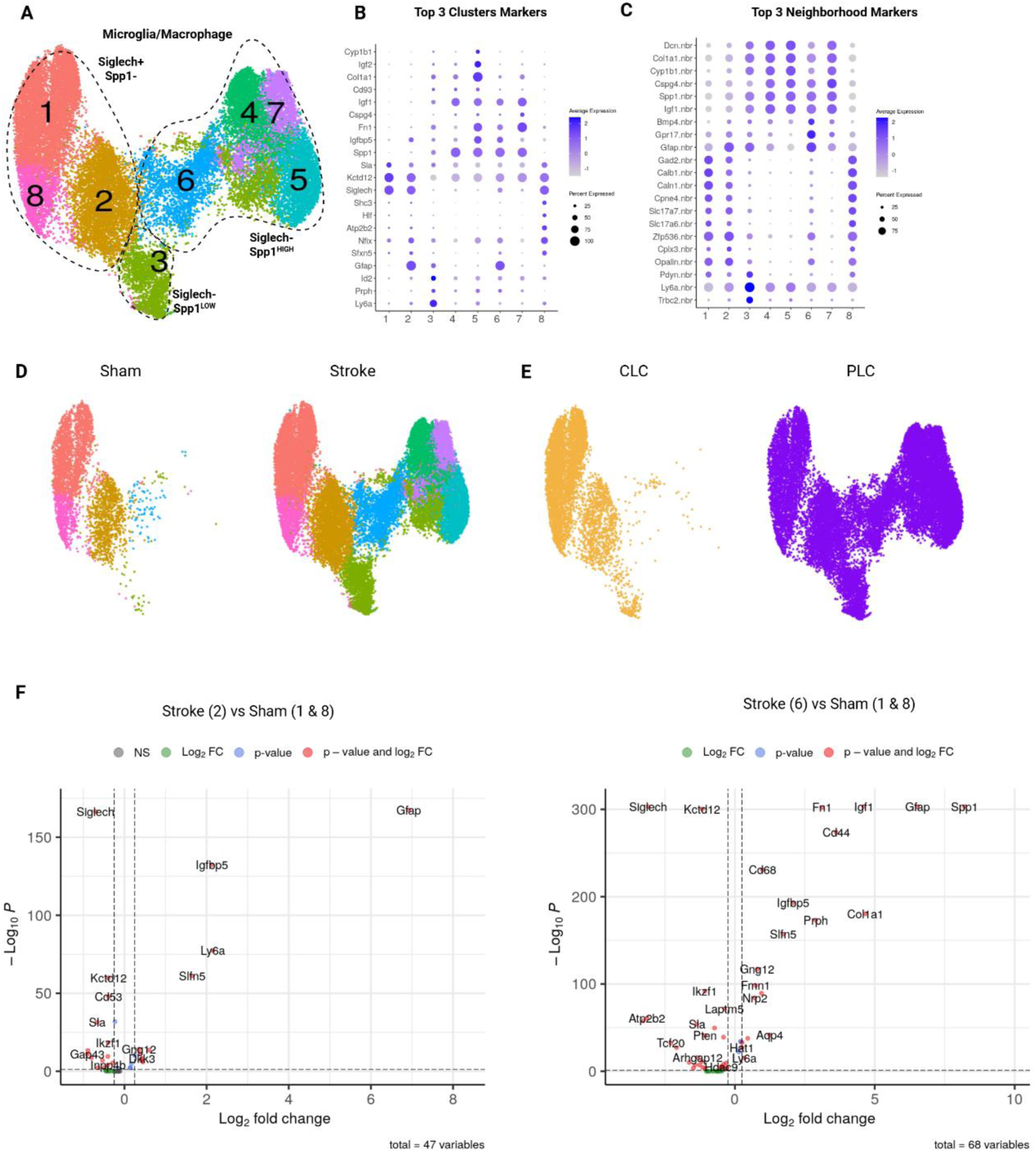
(A) UMAP of BANKSY-clustered myeloid cells annotated for marker genes (B) Top 3 cluster marker dotplot (C) Top 3 BANKSY neighborhood marker dotplot (D,E) UMAPs of myeloid cells split by condition and region (F) Volcano plots of Stroke cluster 2 vs homeostatic populations Sham 1 & 8 (left) and Stroke cluster 6 vs homeostatic populations Sham 1 & 8 (right). Wilcox: |Logfc| > 0.25, adjusted p-value < 0.05.

**Supplemental Figure 4:**
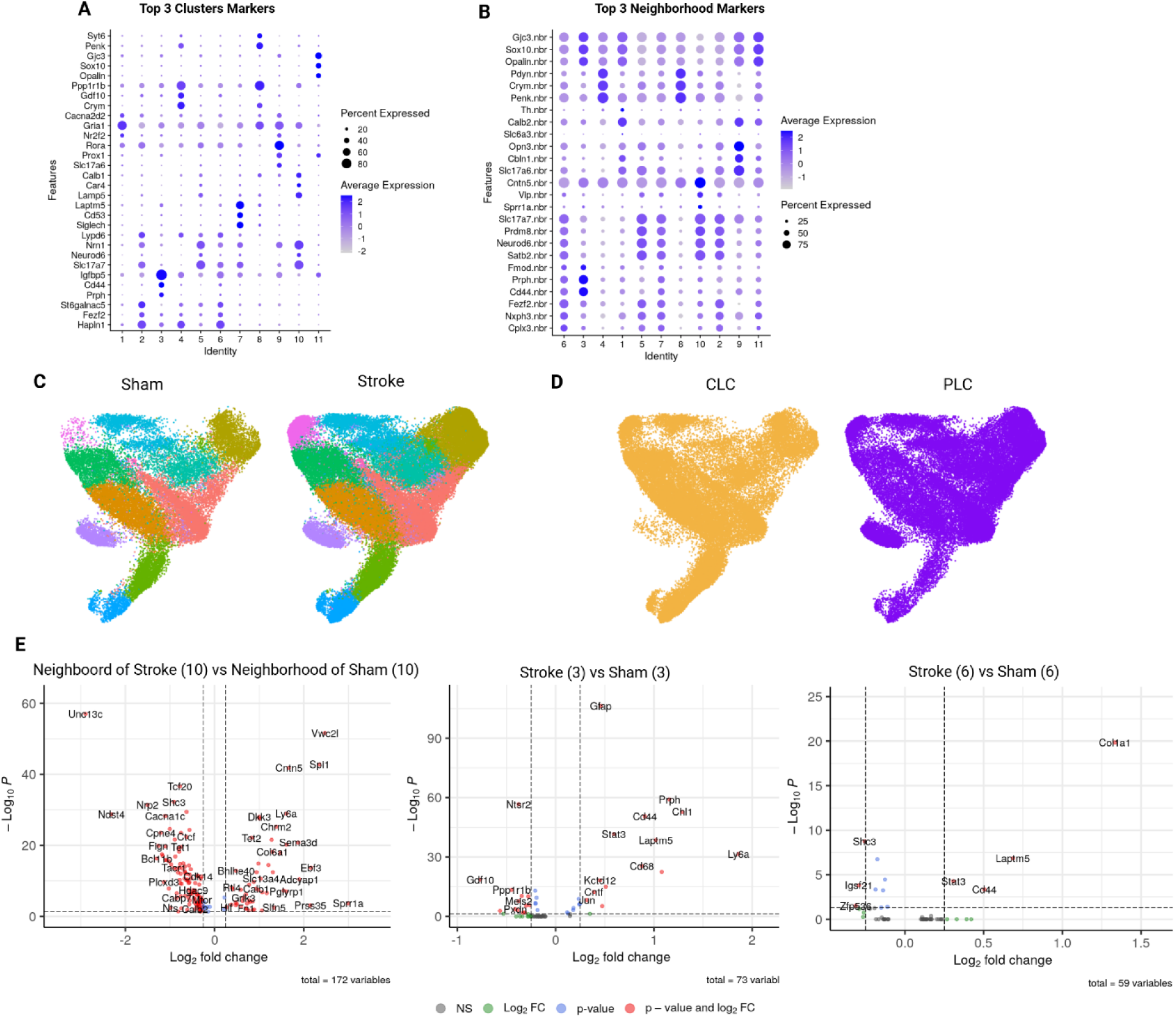
(A) Top 3 cluster marker dotplot (B) Top 3 BANKSY neighborhood marker dotplot (C,D) UMAPs of astrocyte cells split by condition and region (E) Volcano plots of the neighborhood of stroke cluster 10 and neighborhood of sham cluster 10 (left), cluster 3 astrocytes stroke vs sham (middle), and cluster 6 stroke vs sham (right). Wilcox: |Logfc| > 0.25, adjusted p-value < 0.05.

**Supplemental Figure 5.**
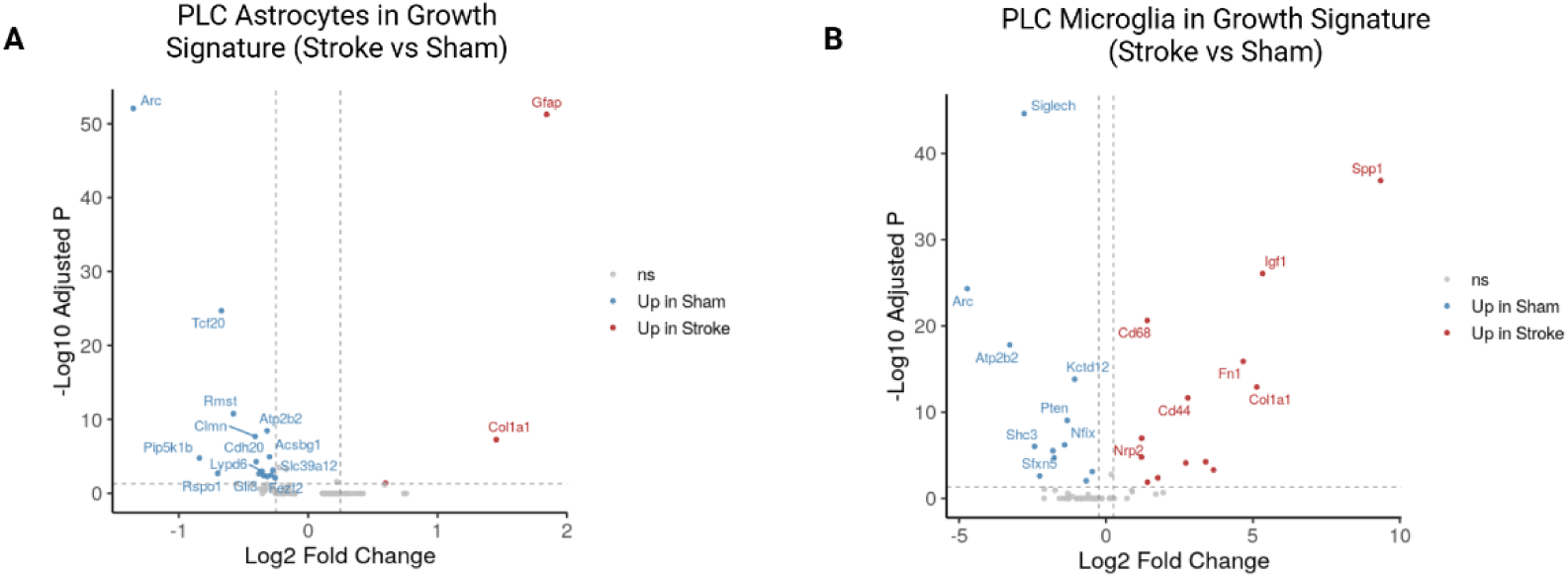
(A,B) Volcano plots of astrocytes (A) and microglia (B) inside the growth signature region of interest stroke versus sham. Logfc > 0.25; adjusted p-value < 0.05.

## Extended Methods

### Animals

C57BL/6J mice (WT) of both sexes were used (12-16 weeks old, 000664 Jackson Laboratory Bar Harbor ME) and maintained on a 12 hour light/dark cycle with standard laboratory chow and water *ad libitum*. All procedures were performed between 9-12am. The Institutional Animal Care and Use Committee of the University of Texas at San Antonio approved all animal procedures.

### Photothrombotic Stroke

To induce focal ischemic strokes, mice were deeply anesthetized with isoflurane and a 45mW 561nm laser (Sapphire 561-FP OEM Laser System, 120mW; Coherent, Santa Clara, CA, USA) is targeted 1.9mm laterally and 0.25mm anterior to bregma to target the right somatomotor cortex. The scalp is shaved, disinfected, and a 1-2cm incision is made to expose the skull. The exposed skull was cleaned with 2% hydrogen peroxide. Rose Bengal (40 mg/kg, Millipore Sigma, 330000-5G) is injected intraperitoneally. Following 1 minute of dwell, the laser is illuminated for 15 minutes. The scalp is then sutured, disinfected, and extended-release buprenorphine is administered subcutaneously. Sham surgeries were performed identically to stroke procedures without illuminating the laser.

### MRI Assessment of Infarct Volumes

All MRIs were acquired on a Bruker 7T system with custom coils at 24 hours post-stroke by the Research Imaging Institute at UT Health San Antonio. Mice were anethesizted by isoflurane and maintained on a breathing and heartrate monitor. Contigious 128x128 px 1mm coronal slices spanning the whole brain AP were collected. We quantified hyperintense lesions by manually tracing them on T2-weighted images in Fiji-ImageJ. Stroke volume was calculated by the sum of outlined lesions across all sections. As this method can be sensitive to acute swelling, we utilized this measurement to normalize relative stroke volumes within groups and across the motor cortex anatomy, not for direct stroke tissue loss.

### Intracranial Injections

For anterograde tracing of the motor cortex, animals underwent identical preparation procedures and were injected with SynaptoTAG4 (AAVDJ-Syn-mTdTomato-Syb2eGFP-WPRE) to label axons with Tdtomato and presynaptic terminals with EGFP. One week prior to stroke, mice received a unilateral AAV2r-SYN-creEGFP into the right motor cortex (-1.8 ML/ 0.0 AP) to label neurons that project to the eventual ischemic lesion. The retrograde labeling was partially bleached by Xenium bleaching protocols and was not utilized in this analysis. AAV interactions with resident immune cells is a known factor and can be seen in this data, however both sham and stroke animals received blinded identical injections to control for this factor.

### Tissue Collection for 10x Xenium

Mice were transcardially perfused with ice-cold RNase-free PBS followed by ice-cold 4% Paraformaldehyde for 5 minutes. Brains were rapidly dissected and placed in 4% PFA for 24-hours followed by 24 hours in 30% sucrose while being maintained at 4°C. Brains were then hemisected, embedded separately in OCT blocks, and stored at -80 °C for long-term storage. Sections were then collected via a cryostat and placed on an ice-cold Xenium slide and stored at -80 °C for 2 weeks.

### 10x Xenium

A 100-gene panel was built on top of the 10x Mouse Brain Panel version 1.1 using the Xenium panel designer (v4.1.0), totaling 347 genes. We processed Xenium slides using the Fixed Frozen protocol with cell segmentation kit staining. Instrument (v4.0.1.4) and analysis (v4.0.1.0). Xenium Ranger outputs of nuclei segmentation using Dapi were utilized to resegment using Proseg ^24^ (v3.1.1) including reissuing Xenium Ranger (v4.0.1.1).

### Xenium Data Processing

Following cell segmentation, data was processed in R (v4.3.2) primarily using Seurat (v5.4.0). Cells were initially filtered for a minimum count of 20. Cell clustering was performed using varying resolutions via Seurat workflows or with BANKSY (k_geom =15, lambda of 0.2,) (v0.1.5). Cell type annotation utilized a multimodal approach. First, we annotated cells using the Allen Brain Institute’s MapMyCells to delineate cell classes and subclasses. Cells with a bootstrap probability of < 0.33 for cell class were removed. PCA clusters with 80% subclass purity were identified as the predominant subclass. For clusters with significant mixed populations, they were first inspected for spatial distribution in the Xenium Explorer and then either subclustered or removed for potential doublets. The immune cell class was subclustered and annotated using a literature-curated single-cell RNA-seq^21,44–46^ dataset of annotated resident and infiltrating immune cells at 7-days post-stroke via gold-standard label transfer in SingleR (v2.4.1). Regions of interest (ROIs) were drawn in Xenium Explorer and exported to R for analysis. Lesion outlines were determined from cell segmentation kit Vimentin staining for the astrocyte border. We manually annotated the cortical surface and cortical-corpus-callosum border to calculate a curved line through the cortex to inform cortical bin segmentation. For growth-associated module scoring, only upregulated growth-associated genes were included and normalized against 10 control genes. As Seurat modules are computed within the dataset as arbitrary units against control gene distribution, we calculated the mean delta between sham (n = 2) and stroke(n = 4) within that bin, allowing for both upregulated and downregulated regions. The same representative sections of CLC and PLC were used from female mice and were chosen for the least tissue artifact (i.e., folding or tearing). Differential expression analysis was conducted using Seurat FindMarkers at a log2 fold-change threshold of 0.25 and a BH-adjusted p-value of < 0.05. UMAPs and clustering were performed on PCs encompassing 90% of the variance. Xenium ranger was used in 10um bins to produce gene density plots.

## Data Availability

Xenium data will be made publicly available upon the publication of the paper and can be requested from the corresponding author.

